# Raman spectroscopy on microcalcifications reveals peculiar differences between male and female breast cancer

**DOI:** 10.1101/2021.01.12.426331

**Authors:** Alessandro Caldarone, Francesca Piccotti, Carlo Morasso, Marta Truffi, Federico Sottotetti, Chiara Guerra, Sara Albasini, Manuela Agozzino, Laura Villani, Fabio Corsi

**Author notes:** Corresponding author: Prof. Fabio Corsi, Department of Biomedical and Clinical Sciences “Luigi Sacco,” University of Milan, Via G. B. Grassi, 74, 20157 Milan, Italy, Surgery Department, Breast Unit, Istituti Clinici Scientifici Maugeri Maugeri S.p.A. SB, via Maugeri 10, 27100 Pavia, Italy. Both authors contributed equally to this work (Alessandro Caldarone, Francesca Piccotti).

## Abstract

Microcalcifications (MCs) are important disease markers for breast cancer. Many studies were conducted on their characterization in female breast cancer (FBC), but no information is available on their composition in male breast cancer (MBC). Raman spectroscopy (RS) is a molecular spectroscopy technique that can explore the biochemical composition of MCs rapidly and without requiring any staining protocol. We here compared, by Raman spectroscopy, the MCs identified on breast cancer pieces from male and female patients. In this study, we used Raman spectroscopy to analyse 41 microcalcifications from 5 invasive MBC patients and 149 MCs from 14 invasive FBC patients. Our findings show that hydroxyapatite is the most abundant type of calcium both in MBC and FBC. However, some differences in the amount and distribution of calcium minerals are present between the two groups. Besides, we observed that MCs in MBC have a higher amount of organic material (collagen) than FBC.

This study provides the first overview of the composition of microcalcifications present in MBC and suggests that they have several specific features. Our result support the need for studies specifically designed to the understanding of MBC.

## Introduction

Male breast cancer is an extremely rare type of malignancy, representing only about 1% of all breast cancer (BC) diagnoses worldwide and only about 1% of cancers occurring in men. [1,2] Despite rare, MBC is associated with higher mortality than female breast cancer. Men are not subjected to screening and thus MBC patients are often older than women and the disease is typically diagnosed at a later stage, with more advanced features. However, even after adjustment for clinical features and access to treatment, a higher proportion of death in males than females was reported. [3] This observation suggests that other factors may underlie such sex disparity and confirms that more studies focusing specifically on MBC are needed.

In fact, since the lack of recruited men in BC clinical trials, and given the relatively few studies on MBC, treatments and diagnostic approaches for men generally derive from those for women. [4] Microcalcifications in MBC are rarer and poorly studied, since mammography is not routinely used on MBC patients, and a mammographic screening for males is not performed. Therefore, MCs in MBC have never been studied in detail, except for their radiological characteristics, since they usually appear rounder and coarser than in FBC. [5] But while many different studies were previously performed on women, we do not have a description of the mineral forms of calcium present in men and we do not know if there are differences in their composition that could highlight biological specificities of MBC.

Raman spectroscopy is a photonic approach that can investigate the composition of samples by irradiation with laser light. RS can provide a complete overview of the mineral constituents of MCs and it is compatible with real-time in vivo diagnostic evaluations. [6] RS was previously used in a few studies for the characterization of MCs in FBC, whereas MBC patients were never included in the studies. [7,8,9,10] In this work, for the first time, we applied RS for the detailed characterization of MCs present in MBC and we compared their composition with a dataset of MCs present in FBC.

## Materials and methods

This study includes 19 patients affected by luminal invasive breast carcinoma and treated at the Breast Unit of the ICS Maugeri in Pavia, a EUSOMA-accredited, tertiary referral center involved in wide screening programs in Northern Italy, between 2013 and 2019 (5 MBC and 14 FBC). From surgical specimens of MBC patients and biopsy specimens of FBC patients, MCs were identified and analyzed on RS (see Supplementary Data for details on data acquisition and analysis). All patients had to sign an informed consent authorized by the Ethics Commission of the Institutions (Protocol 2281/2018 EC), which approved the study in accordance with the Helsinki Declaration. Only patients with a diagnosis of invasive ductal carcinoma of the breast were included.

## Results

Detailed patient characteristics are shown in Table S1-S2 in the supplementary information. 190 MCs were identified and fully scanned by RS: 41 MCs from 5 MBC patients and 149 MCs from 14 FBC [7]. By the RS analysis of MCs, some differences in the proportion of minerals and their crystal characteristics emerged.

Hydroxyapatite (HA), a particular form of calcium phosphate, appeared as the most common mineral present in both groups of microcalcifications; in particular, HA was identified in 40 male MCs (Figure 1a) and 136 female MCs (Figure 1c). Overall, HA made 92% of the total calcium deposit found in MBC (Figure 1b) and 96% of the one in FBC (Figure 1d). A different form of calcium phosphate named whitlockite (WIT) was detected in 2 male MCs (Figure 1a) and 6 female MCs (Figure 1c), always in association with HA. WIT resulted more abundant in MBC (3,1 %) than in FBC (0,35 %) (Figure 1b-1d).

**Fig. 1.**
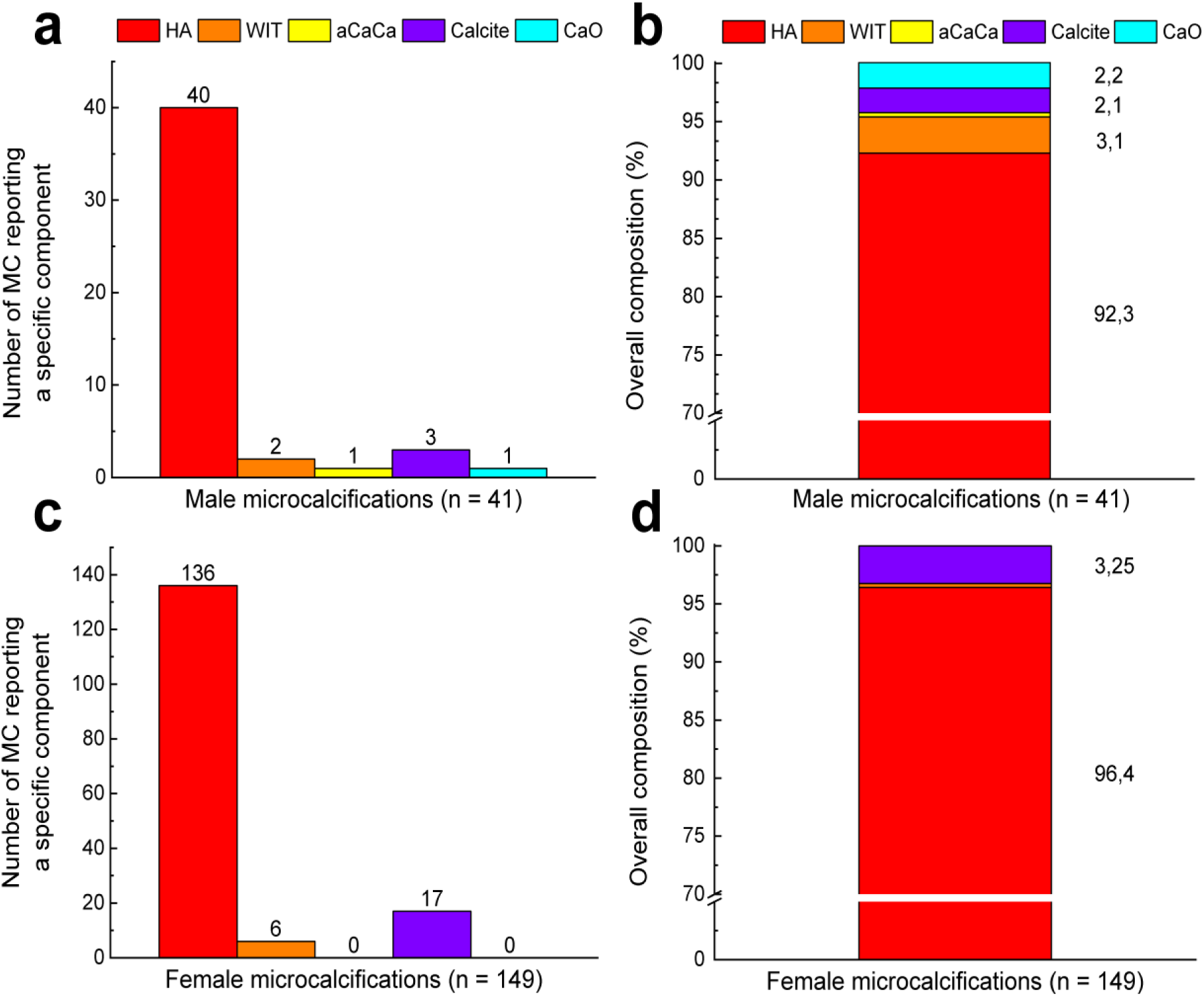
Detailed composition of representative microcalcifications. Number of male **a)** and female **c)** microcalcifications exhibiting at least one pixel of the mineral components described by the legend. Overall composition of male **b)** and female **d)** microcalcifications, calculated, considering altogether all microcalcifications

Calcium carbonate is present in MCs from both MBC and FBC in its crystalline form (calcite). However, while in FBC calcite was always detected as a small deposit within larger MCs made of HA, in one MBC patient we identified a MC entirely made of calcite with a small deposit of amorphous calcium carbonate (aCaCa) inside. At last, one male MC resulted made of calcium oxalate (CaO) in combination with a smaller deposit of HA and WIT (Table 1a). This represented an anomalous result as CaO was not detected in our dataset of MCs from FBC (Table 1b) and, to the best of our knowledge, was never reported in the scientific literature in infiltrating FBC.

**Table 1.** Number of male a) and female b) microcalcifications exhibiting at least one pixel of the mineral components described by the legend

| **a** |  |  |  |  |  | **b** |  |  |  |  |  |
| --- | --- | --- | --- | --- | --- | --- | --- | --- | --- | --- | --- |
|  | HA | WIT | aCaCa | Calcite | CaO |  | HA | WIT | aCaCa | Calcite | CaO |
| **HA** | 40 | 2 | 0 | 2 | 1 | **HA** | 136 | 6 | 0 | 4 | 0 |
| **WIT** |  | 2 | 0 | 0 | 1 | **WIT** |  | 6 | 0 | 0 | 0 |
| **aCaCa** |  |  | 1 | 1 | 0 | **aCaCa** |  |  | 0 | 0 | 0 |
| **Calcite** |  |  |  | 3 | 0 | **Calcite** |  |  |  | 17 | 0 |
| **CaO** |  |  |  |  | 1 | **CaO** |  |  |  |  | 0 |

We also compared the crystal characteristics of calcium phosphate deposits identified by RS in the two groups of MCs.

The analysis of Raman spectra confirmed that MCs present MBC and FBC share a similar profile. However, the different spectrum (Figure S1) highlighted small differences in the crystal structure between the two groups. A detailed analysis revealed the significance of the spectral differences (Figure S2). In particular, the bands at 960 cm-1, relative to the presence of phosphate groups, and the one at 1075 cm-1, relative to carbonate, are significantly reduced in MBC than in FBC (Figure 2a-2b). On the contrary, the band at 1462 cm-1 relative to the presence of organic material within MCs (mostly referring to proteins [11]), is more pronounced in MBC (Figure 2c). We also studied the ratio between the bands at 960 cm-1 and 1073 cm-1 that, on the contrary, did not vary between MBC and FBC (Figure 2d), suggesting that the difference in the crystal structure is driven by a different amount of proteins and other organic materials present within the MCs.

**Fig. 2.**
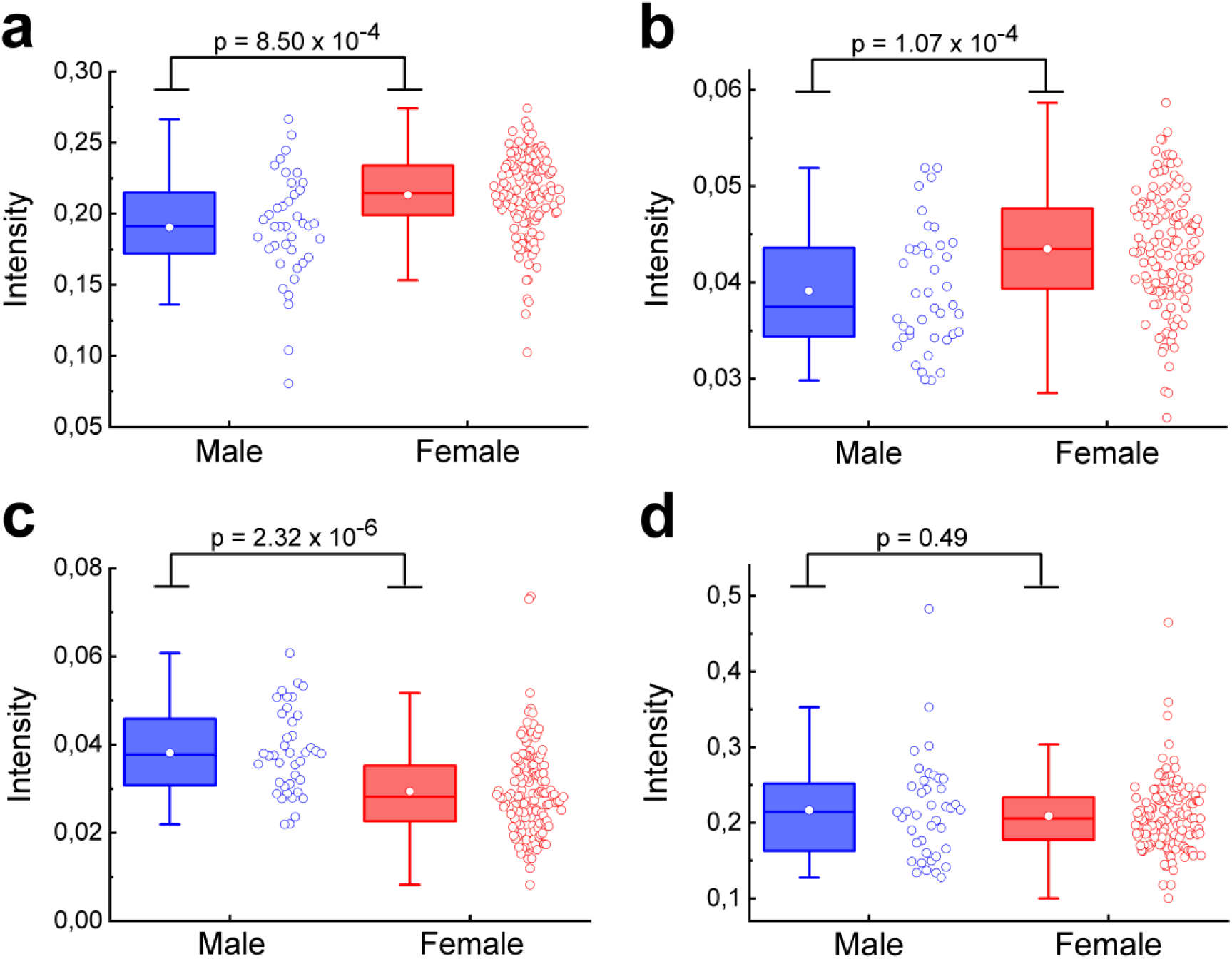
**a)** Box plot reporting the intensity of the phosphate band (960 cm^-1^) in MBC and FBC. **b)** Box plot reporting the intensity of the carbonate band (1075 cm^-1^) in MBC and FBC. **c)** Box plot reporting the intensity of the band relative to the organic material (1462 cm^-1^) in MBC and FBC. **d)** Ratio between the band of carbonate and phosphate in MBC and FBC

## Discussion

The role of MCs in BC is still not fully understood. However, their association with the presence of a malignant lesion is well known and MCs are an important marker of the disease. A recent study by our group proved that the mineral characteristics of MCs correlate with the histological classification of FBC making the connection between MCs and BC even tighter [7].

By using RS imaging, we now studied the different forms of calcium present in MCs from infiltrating MBC and we compared the results with the cohort of infiltrating FBC included in our previous study.

The results show that MCs in MBC resulted mainly made of HA, as previously reported also for FBC. However, some clear differences emerged during the analysis. WIT, a different mineral form of calcium phosphate is a not negligible component in MBC while was present just as traces in infiltrating FBC. In one MBC, we also identified the presence of CaO. This is particularly notable, as CaO is a specific marker of benign lesions and was never reported on invasive FBC. [12,13] Besides, in FBC CaO forms homogeneous calcification distinct from the one made of calcium phosphate. [14] On the contrary, here CaO co-localized with a deposit of HA and WIT. [15] Figure S2 in supporting information report the MC presenting both calcium oxalate and calcium phosphate present as HA and WIT.

The comparison of the spectral characteristics between MBC and FBC also highlighted differences in the crystal structure of MCs. MCs in MBC are richer in proteins and have less calcium. However, the ratio between the peaks of phosphate and of carbonate inclusion is unchanged between the two groups. These data suggest an active role of collagen, and thus of cancer microenvironment, on the formation of MCs and support the fact that MBC and FBC are diseases with proper characteristics that should be taken into the account.

## Conclusion

The bio-mineralization process that led to the formation of MCs in BC is currently under investigation by several groups as MCs are recognized as an early marker of BC. The present study reports the first detailed characterization of MCs present in MBC. These findings highlighted some particular features of MCs in MBC in terms of the different mineral forms of calcium and in the balance between calcium and proteins. Overall, our data suggest that different biochemical processes can influence the formation of MCs in MBC an FBC; and support the need to design studies for the understanding of MBC.

## Supplementary

Supplementary file “ESM.pdf” contains details about the clinical cohort involved in the study and an accurate description of the experimental protocols.

## Declarations

### Ethics approval

All procedures performed in this study involving human participants were in accordance with the ethical standards of the Ethics Commission of the Institutions (Protocol 2281/2018 EC), which approved the study in accordance with the Helsinki Declaration.

This article does not contain any studies with animals performed by any of the authors.

### Consent to participate

Informed consent was obtained from all individual participants included in the study.

### Data availability

The data generated and analysed during the current study is available from the corresponding author on reasonable request.

## Authors’ contribution

C. Morasso, F. Corsi: study concepts; C. Morasso, F. Corsi, L. Villani: study design; A. Caldarone, F. Piccotti, C. Guerra, M. Agozzino: data acquisition; A. Caldarone: quality control of data and algorithms; C. Morasso, F. Corsi, F. Sottotetti: data analysis and interpretation; S. Albasini: statistical analysis; C. Morasso, A. Caldarone, F. Piccotti: manuscript preparation; F. Sottotetti: manuscript editing; F. Corsi: manuscript review.

