## Supplementary material for "Raman spectroscopy on microcalcifications reveals peculiar differences between male and female breast cancer"

*Corresponding author:*

### Subjects included:

The study included five male and fourteen female breast cancer patients treated at the Breast Unit of the ICS Maugeri in Pavia between 2013 and 2019, on which microcalcifications were identified on the collected tumor pieces. In order to be included in the study, all patients had to sign an informed consent authorized by the Ethics Commission of the Institutions (Protocol 2281/2018 EC), which approved the study in accordance with the Helsinki Declaration.

The age of the patients at the time when they were diagnosed with carcinoma varies between 50 and 76 years, and they all have the same diagnosis: "luminal invasive breast carcinoma". Detailed male and female patients characteristics are shown in Table S1 and Table S2.

**Table S1 Description of the male patients database included in the study. For each patient is reported the age at the time when he was diagnosed with carcinoma, the number of microcalcifications identified on the collected tumor pieces, the molecular subtype of the carcinoma and its diagnosis. Abbreviations: IDC, invasive ductal carcinoma.**

| Samples | Patient's age at diagnosis | Microcalcifications identified | Molecular subtype | Diagnosis |
|---|---|---|---|---|
| MBC1 | 62 | 15 | Luminal A | IDC, G2 SBR |
| MBC2 | 76 | 11 | Luminal B | IDC, G2 SBR |
| MBC3 | 75 | 5 | Luminal B | IDC, G2 SBR |
| MBC4 | 71 | 5 | Luminal A | IDC, G2 SBR |
| MBC5 | 76 | 5 | Luminal B | IDC, G2 SBR |

**Table S2 Description of the female patients database included in the study. For each patient is reported the age at the time when he was diagnosed with carcinoma, the number of microcalcifications identified on collected tumor pieces, the molecular subtype of the carcinoma and its diagnosis. Abbreviations: IDC, invasive ductal carcinoma; ILC, invasive lobular carcinoma; IMC, invasive mucinous carcinoma.**

| Samples | Patient's age at diagnosis | Microcalcifications identified | Molecular subtype | Diagnosis |
|---|---|---|---|---|
| FBC1 | 71 | 2 | Luminal B | IDC, GX SBR |
| FBC2 | 70 | 2 | Luminal B | IDC, G3 SBR |
| FBC3 | 50 | 21 | Luminal A | IDC, G2 SBR |
| FBC4 | 75 | 12 | Luminal B | IDC, G2 SBR |
| FBC5 | 69 | 3 | Luminal B | IDC, G2 SBR |
| FBC6 | 66 | 2 | Luminal A | ILC, GX SBR |
| FBC7 | 65 | 18 | Luminal A | IDC, G2 SBR |
| FBC8 | 73 | 25 | Luminal A | IDC, G2 SBR |
| FBC9 | 59 | 28 | Luminal A | IDC, G2 SBR |
| FBC10 | 72 | 1 | Luminal B | IDC, G2 SBR |
| FBC11 | 50 | 12 | Luminal B | ILC, G1 SBR |
| FBC12 | 54 | 5 | Luminal A | IDC, G2 SBR |
| FBC13 | 55 | 5 | Luminal A | IDC, G2 SBR |
| FBC14 | 73 | 7 | Luminal A | IMC, GX SBR |

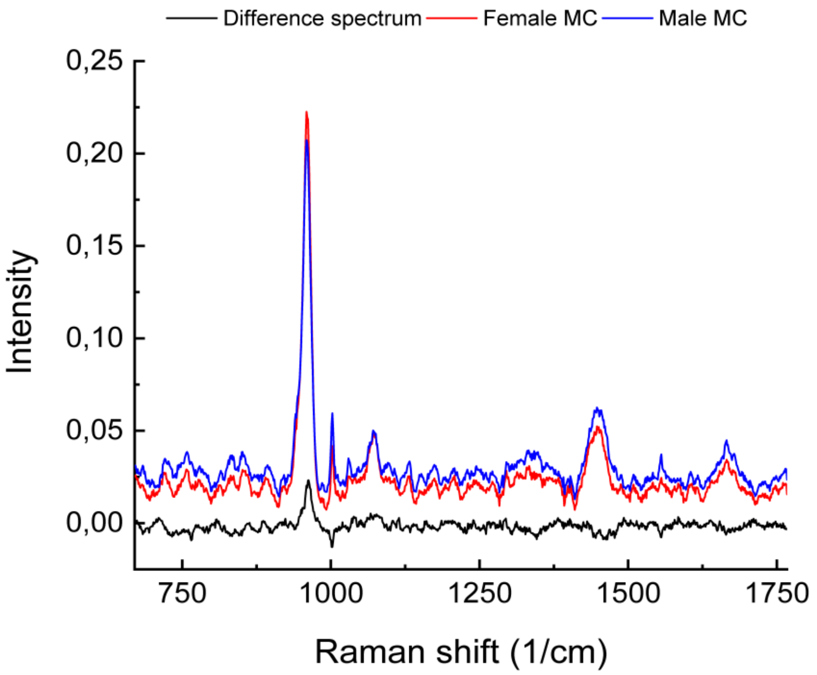


**Fig. S1** Mean spectra of MCs found on MBC (blue) and FBC (red). Black line is the differential spectrum between the two categories (FBC-MBC)

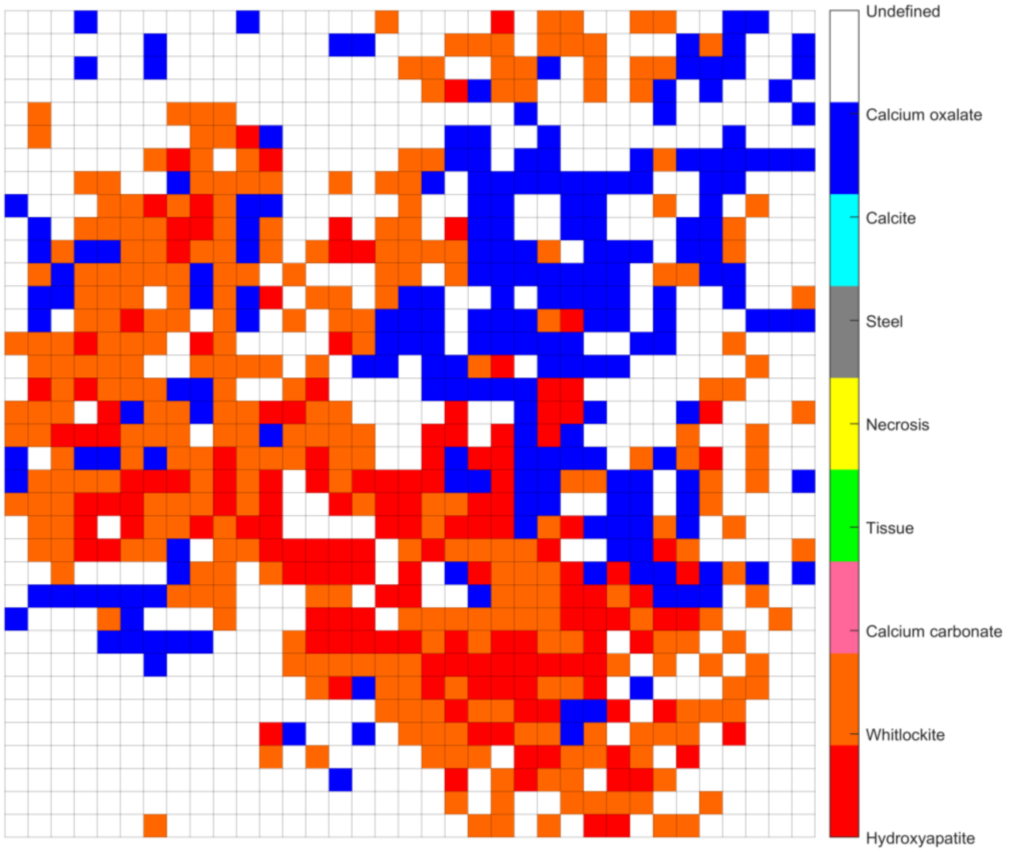


**Fig. S2** Raman map illustrating a microcalcification from MBC where Hydroxyapatite, Whitlokite and Calcium Oxalate were found in close connection

### Detailed experimental protocols:

#### Tissue preparation

Tissue slices for Raman analysis were prepared according a previously published protocol (Vanna e C). Briefly, tissue slices were generated from formalin-fixed, paraffin embedded tissue blocks. For each patient, a 10-mm slice was microtomed and mounted on mirrored stainless steel slides specific for Raman measurements (Renishaw plc). The 10-mm slices selected for Raman analyses were then dewaxed by two baths of hexane 95% (Merck KGaA), two baths of ethanol absolute (Merck KGaA), and a final bath of ethanol 95%, followed by air drying for 2 hours.

#### Raman Spectroscopy

A Renishaw InVia Reflex Confocal Raman Microscope was used to acquire the Raman maps. The measurements were carried out with a 785 nm laser source, with power set at 50 mW, coupled with a Leica 100x N-Plan lens; with grating and center set at 1200 I/mm and 1250 $cm^{-1}$ respectively, spectra with Raman shift between 670-1767 $cm^{-1}$ (1015 points) were obtained. The detector present in the spectrometer is a CCD (Charge Coupled Device) (1024 * 256 pixels) sensitive between 400 and 1060 nm and is cooled to a temperature of -70°C.

Alignment and calibration of the instrument are two procedures that are performed daily using the WiRe (Renishaw) software.

On each microcalcification a square Raman map with steps between 4 and 15 μm, depending on the size of the microcalcification, and raster direction has been acquired; thus, each map contains between 1000 and 4000 spectra. Finally, the acquisition time of a single spectrum has been set to 3 seconds with a single repetition.

#### Data processing of Raman data

Raman data were acquired using the WiRe softaware from the producer of the instrument (Renishaw). MATLAB (MathWorks) and OriginPro2019 (Originlab Corporation) were used for preprocessing and data analysis.

The first pre-processing operation carried out on the Raman data was the removal of cosmic rays. To perform this task it was used the CRR (Cosmic Ray Remover) options of the WiRE software, based on the nearest neighbor algorithm.

Spectra were denoised using a Singular Value Decomposition (SVD), a matrix factoring technique based on the use of eigenvalues and cars. Baseline signals referring to florescence of the sample were removed using an Asymmetric Least Squares (ALS) approach and the obtained spectra were vector normalized. SVD, baseline correction and normalization operations were performed on MATLAB.

After data pre-processing operations each spectrum was assigned to one of the pre-selected components: Hydroxyapatite, Whitlockite, Calcium Carbonate, Calcite, Calcium Oxalate, Tissue,

Steel and Necrosis. The first five components are the mineral components most present in Type I and Type II microcalcifications; the last three, on the other hand, although not of particular interest in this study, were added because some of the acquired spectra showed their typical spectral characteristics.

The criterion used to carry out this type of assignment is the Pearson correlation index (also called linear correlation coefficient); so, for each spectrum of each Raman map the correlation with each of the reference spectra of the components was calculated and then, on the basis of the results obtained, the spectral classification was carried out. A correlation coefficient value equal to or greater than 0.3 was chosen as criterion for the assignment of each pixel to one of the component. Spectra with correlation coefficient minor than 0.3 were considered too noisy to be accurately attributed to one category and were classified as "Not defined". Furthermore, for spectra classified as tissue (according to Pearson index), is the result is confirmed or modified using two other indices: peaks numbers, and peaks positions. For each of these tissue spectra the algorithm calculates these parameters and compares them with the number and position of the peaks of the eight reference spectra; finally, the three indices vote and the final classification is obtained for these spectra which were initially assigned as tissue. Thus, for each Raman map, a false-colour image was generated that contains a classification based on only spectral information.

The reference spectra used for the classification were produced by acquiring Raman signals from real samples and confirming their nature through the analysis of X-ray scattering and/or through data in the literature. As far as hydroxyapatite and whitlockite are concerned, their nature has been confirmed both with the data present in the literature and with the X-ray scattering analysis on adjacent 50 μm slices from the same sample; for calcium carbonate, calcite and calcium oxalate, the small number and size of the microcalcifications in which these components were found did not allow to confirm their nature through the analysis of the scattering, so we had to limit ourselves to the data present in the literature.

After the data processing and classification, a quantitative analysis were carried out to study the general composition of the microcalcifications incorporated in the study and to understand which components were the most identified.

In order to make a comparison between the microcalcifications found on tumor pieces removed from the breast of male patients and the microcalcifications identified on biopsies from female patients, a single spectrum was considered for each map, calculated by averaging the spectra classified as hydroxyapatite and whitlockite; this decision was taken because most of the microcalcifications analyzed are Type II, i.e. they are mainly composed of calcium phosphate. Raman maps in which these components were not identified were not considered for this analysis.

Continuous variables were compared using the non-parametric t-Student test for variables with normal distribution, and with a level of statistical significance set to $P<0.05$.
